# Whole group tracking reveals that relatedness drives consistent subgrouping patterns in white-nosed coatis

**DOI:** 10.1101/2023.12.14.571650

**Authors:** Emily M. Grout, Josué Ortega, Pranav Minasandra, Matthew J. Quin, Margaret C. Crofoot, Ariana Strandburg-Peshkin, Ben T. Hirsch

## Abstract

The formation of subgroups can allow group-living animals flexibility to balance the costs and benefits of sociality over time. Subgrouping dynamics emerge from individual decisions about whether and with whom to maintain cohesion, with these decisions potentially influenced by ecological, physiological, and social factors. We GPS-tracked the movements of nearly all members of three wild white-nosed coati (*Nasua narica*) social groups that differed in their demographic profiles to better understand how these highly social, frugivorous carnivores weight the relative importance of these different factors in their grouping decisions. Quantifying group movements and subgrouping patterns, we found that two of the three groups we tracked exhibited fission-fusion behaviours, with groups splitting into subgroups that persisted over varying timespans from minutes to days. In contrast, the third group remained together across the entire observation period. When groups split, they did not do so randomly; instead, individuals tended to form subgroups with the same individuals consistently over time. Assessing the drivers of subgrouping patterns revealed that subgroup membership was associated with genetic relatedness, but not physiological similarity as quantified by age and sex homophily. Our results demonstrate great variation in subgrouping patterns within a single species, while also highlighting a consistent role of relatedness in driving social preferences when subgroups form.

## Introduction

Social animals are strongly influenced by their neighbours; thus, the size, structure, and demography of their groups are important drivers of individual behaviour and fitness. When groups forage cohesively, individuals must balance resource competition, which can drive individuals apart, with predation avoidance, which generally bring animals together (Hirsch, 2007). To balance the costs and benefits of grouping, some animals flexibly adjust their patterns of spatial association, breaking into smaller sub-groups or fusing into larger aggregations in response to local conditions. The spatiotemporal scale and social boundaries of these fission-fusion dynamics vary substantially among species, from bird flocks and fish schools where individuals join and leave frequently and freely to the more constrained splitting and merging within defined social communities seen in chimpanzees, guineafowl, and hyaenas (Holekamp et al., 1997; Kelley et al., 2011; Mcfarland Symington, 1990; Papageorgiou et al., 2019; M. J. Silk et al., 2014). Because fission-fusion dynamics emerge from sets of individual decisions about the relative costs and benefits of association, they provide insight into the social and environmental conditions that favour the formation, maintenance and, ultimately, the evolution of animal societies.

Ecological, social, and physiological factors are all known to influence the formation and composition of subgroups in species that show fission-fusion dynamics (Aguilar-Melo et al., 2020; Aureli et al., 2008; Grueter et al., 2023). While ecological factors typically drive overall subgrouping tendencies (Aureli et al., 2008; Sueur et al., 2011; van Schaik & Brockman, 2009), the composition of the resulting subgroups is often influenced by the characteristics and relationships among group members (Sueur et al., 2011). Variation in physiological traits, for example, can influence subgroup membership when individuals have different needs or physical constraints that cause them to split apart (Conradt & Roper, 2000; Hartwell et al., 2014; Krause et al., 2002; Matthews et al., 2021). In such cases the resulting subgroups are expected to be composed of individuals with similar physiological traits (Clutton-Brock et al., 1977; Ruckstuhl, 1998). Social factors, such as the strength of social bonds, position in a dominance hierarchy, and kinship, can also shape subgroup composition, with group splits allowing individuals to avoid group members from whom they receive aggression, while remaining with those who tolerate them or from whom they receive benefits (Carter et al., 2013; M. J. Silk et al., 2014; Sueur et al., 2011). While both physiological and social factors can be important for driving subgrouping patterns, these factors may act in opposition. When kinship drives association due to differences among related individuals in age or sex, substantial physiological heterogeneity within subgroups is expected to emerge. Conversely, individuals with similar preferences and constraints may not be related. Patterns of fission-fusion behaviour can thus shed light on the main drivers of social cohesion in a given species.

Fission-fusion dynamics can provide insights into the costs and benefits of grouping, and have been implicated in the evolution of complex sociality and large brain size (Aureli et al., 2008; Holekamp et al., 2007). Unfortunately, quantifying how groups split and merge in order to understand the decision-making processes that drive these patterns presents a methodological challenge. It is practically infeasible for human observers to simultaneously record the location and behaviour of all members of social groups, especially when they have split into multiple subgroups. Field studies that employ direct observation typically locate or track one subgroup at a time, leaving the movement patterns of other group members unrecorded (Grueter et al., 2023; Hartwell et al., 2021). Given that the frequency of fission-fusion events and the composition of subgroups may vary across the landscape, traditional methods may lead to biases in which subgroups and events are observed. Tracking technologies such as GPS tags offer the potential to monitor the movements of multiple individuals simultaneously (Della Libera et al., 2023; Kays et al., 2015; Strandburg-Peshkin et al., 2015). This approach can give us greater insight into the decision-making processes of group members by allowing us to determine which subgroups individuals choose to join as well as those they reject.

White-nosed coatis (*Nasua narica*) live in stable heterogeneous social groups that can break up into smaller foraging parties during the day (Gilbert, 1973; Gompper, 1997; Kaufmann, 1962; Romero & Aureli, 2007). They are generalist, opportunistic foragers that predominantly feed on fruit and leaf litter invertebrates (Gompper, 1996b; Hirsch, 2009; Valenzuela, 1998). In response to reduced fruit availability coatis forage in smaller subgroups (Gompper, 1996a, 1997). Their social and ecological flexibilities enable them to thrive across a broad spectrum of forested habitats spanning Central, South, and North America (Frey et al., 2013; Nigenda-Morales et al., 2019; Valenzuela & Ceballos, 2000). Groups range in size from four to over 30 individuals and typically consist of multiple adult females and their dependent offspring (Gompper, 1996a; Hirsch & Gompper, 2018; Kaufmann, 1962). These groups are primarily composed of highly related individuals; however, they may contain unrelated females which often receive a disproportionate amount of aggression from other group members (Gompper et al., 1997). Adult males are predominantly solitary, except during the breeding season when they often temporarily join female groups (but see Gompper & Krinsley, 1992). Predation risk can play an important role in spatial structure of coati groups, with juveniles, who are at the highest risk of predation due to their smaller body size, positioned in close proximity to one another (Hirsch & Gompper, 2018; Russell, 1979). Despite our understanding of the spatial structure of coati social groups, little is known about how groups dynamically change composition.

Here, we use simultaneous tracking of entire groups to quantify coati fission-fusion dynamics, including the frequency of splits and merges, the distribution of subgroup sizes and patterns of subgroup membership. With data from three groups that varied in their demographic and relatedness structures, we test whether individuals show consistent subgrouping patterns across time, and the extent to which group splits are driven by social vs physiological factors. If social bond strength drives subgrouping patterns, we predict that group fissions will take place along kinship lines (Gompper et al., 1997). If physiological factors drive fission-fusion dynamics, we expect subgroups to divide according to age/sex class (Harel et al., 2021).

## Methods

### Study site and data collection

Fieldwork was conducted in Soberania National Park (SNP) and on Barro Colorado Island (BCI), Panama. Both study sites consist of semi-deciduous lowland tropical forest. Although the sites are only 5 km apart, they have been isolated from one another since 1914 when the Chagres River was dammed to create Lake Gatun and the Panama Canal. We equipped three groups of wild white-nosed coatis (*Nasua narica*) with custom built collars that recorded group members’ positions and activity patterns using GPS and IMU sensors (e-Obs Digital Telemetry, Gruenwald, Germany) as well as their vocalisations using audio recorders (Soroka 18E, TS-Market). Only the GPS data were used in this study. Coatis were caught using Tomahawk traps and chemically immobilised using Telazol (50 mg/mL tiletamine and 50 mg/mL zolazepam; 5.4 ± 0.5 mgs/kgs). For two of the three groups (Trago and Presidente), all members were collared. In the third group (Galaxy), we collared all members except one adult female (10/11 or 92% of group members). Members of the Galaxy group were collared during the mating season, which is when adult males temporarily join groups, therefore we also collared the adult male associating with the group during this time (see Table 1 for details on group composition and tracking times).

**Table 1.** Tracking periods and group composition with age/sex class composition for the three coati study groups collared in Soberania National Park (SNP) and on Barro Colorado Island (BCI), Panama. Adult males are generally solitary and therefore were excluded from the group collared and group size values.

| Group | Tracking period | Location | Adult |  | Sub-adult |  | Juvenile |  | group collared/<br>group size |
| --- | --- | --- | --- | --- | --- | --- | --- | --- | --- |
|  |  |  | ♀ | ♂ | ♀ | ♂ | ♀ | ♂ |  |
| Galaxy | 24.12.2021 – 13.01.2022 | SNP | 7 | 1 | 1 | 2 | 1 | 0 | 10/11 |
| Trago | 24.03.2022 – 10.04.2022 | SNP | 0 | 0 | 0 | 1 | 5 | 1 | 7/7 |
| Presidente | 19.01.2023 – 02.02.2023 | BCI | 3 | 0 | 5 | 2 | 2 | 4 | 16/16 |

GPS data were recorded at a rate of one fix per second from 0600 to 0900 each day. From 0900 to 1800, a burst of six GPS points was recorded every 10 minutes. During the night (1800 to 0600) a burst of six GPS points was recorded every hour. The mean GPS fix success rate was 96.5%, 96.1%, and 96.9% for Galaxy, Trago, and Presidente groups respectively. The average relative GPS error (measured under dense canopy typical of the coatis’ habitat as the relative error between two collars at known distances apart) was 3.86 ± 1.06 m. For collar recovery, automated drop-off devices (Micro-TRD, Lotek) were incorporated into the collars, and programmed to activate on day 18 of collar deployment. Due to a technical problem, all drop-offs from the first group (Galaxy) failed to activate. For the two subsequent groups, this issue was resolved, resulting in 78% and 100% drop-off success rates for the Trago and Presidente groups respectively. For cases where drop-offs failed, coatis were recaptured and all collars were successfully removed within four weeks of the drop-off activation date.

### SNP genotyping and relatedness estimates

We used SNP genotyping of tissue samples collected during captures to calculate relatedness between members of the coati groups. While collaring individuals, we collected tissue samples from all study individuals and stored them in 95% ethanol for genetic analysis. We submitted the tissue samples to the University of Minnesota Genomics Center (UMGC) for genotyping-by sequencing (GBS). High quality genomic DNA were extracted from samples using a QIAGEN DNeasy Blood and Tissue Kit following manufacturers protocols, and a double digest restriction-site associated DNA method (ddRAD) was followed, using the *BamHI* and *Nsil* restriction enzymes for digestion. Illumina primers and individual barcodes were ligated to DNA fragments and amplified using PCR. Sequencing was performed using the Illumina NextSeq 2000 platform with a 100-cycle run configuration, resulting in approximately 6.5 million reads per sample.

We used FastQC v0.11.9 software (Andrews, 2010) to assess and analyse the quality of the raw sequence reads. We trimmed adapter and padding sequences using a custom UMGC perl script gbstrim.pl (Garbe, 2023), and aligned to a *Nasua narica* reference genome using the Burrows-Wheeler Aligner tool in BWA v0.7.17 (Li & Durbin, 2009). The aligned sequence files were sorted and indexed using SAMtools v1.6 (Li et al., 2009), and variants across all samples were called using FreeBayes v1.2.0 (Garrison & Marth, 2012) with the parameters: –use-best-n-alleles 4 –min-coverage 102 –limit-coverage 500. We removed low quality variants using the vcffilter tool in vcflib v1.0.1 (Garrison et al., 2022), specifying the option: -f QUAL > 20. We removed any samples containing more than 50% missing genotypes, and any variants with genotype calls in less than 95% of samples, as were variants with a minor allele frequency of less than 1%.

We converted the filtered variant file containing 37 samples and 10,972 Single Nucleotide Polymorphisms (SNPs) into a compatible file for input into COANCESTRY v.1.0.1.10 (Wang, 2011) using the *dartR* v2.9.7 package (Mijangos et al., 2022) in R (R Core Team, 2017). Although a variety of relatedness estimators exist for analysing SNP data, we implemented the triadic maximum likelihood estimator (TrioML; Wang, 2007) to calculate relatedness coefficients among each unique coati pair. We made this decision as each of the seven estimators available in COANCESTRY software were highly correlated for the coati dataset (r > 0.93 for each comparison between relatedness estimators), and hence the TrioML estimator was selected as the most suitable option as it is also considered robust to inbreeding, small sampling sizes, and genotyping error (Hauser et al., 2022; Wang, 2007).

### Ethical Note

All methods of data collection were performed in accordance with the Institutional Animal Care and Use Committees guidelines. Smithsonian ACUC clearance number (2017-0815-2020). Biological samples export permit number (PA-01-ARG-160-2022). Methods followed the appropriate ethical guidelines set by the American Society of Mammologists.

### Analyses

#### GPS data processing

We used RStudio (R Core Team, 2017) for all analyses. We downsampled the 1 Hz GPS data (recorded from 0600 – 0900) to one GPS point every 10 minutes, which resulted in a consistent sampling interval of 1 fix every 10 minutes across the entire active period (0600 – 1800). One of the Galaxy group members (Venus) remained stationary in a tree for three days after collaring before re-joining the group, which may have been a response to the capture. These data points were removed from the analysis. Two group members from Presidente group (Peron and Moscoso) wore collars which fell off before the scheduled drop-off date, and these individuals were refitted with new collars. The data recorded from the fallen collars were removed from the analysis, and this resulted in a gap of 101 hours for Peron and a gap of 5 hours for Moscoso.

#### Identifying subgroups

We defined subgroup membership at each time step using density-based spatial clustering (DBSCAN) (Ester et al., 1996). This algorithm is similar to the “chain rule” employed in previous observational field studies to quantify group membership (Whitehead, 2008), where a group is defined as a set of individuals who are within a distance *ε* of the nearest group member. We set the noise parameter of the DBSCAN algorithm to 1, meaning that groups of all sizes were identified. We ran the DBSCAN analysis at a range of spatial scales and the results remained qualitatively similar but with more groups identified for shorter values of *ε* as expected (see S1 for results at different *ε-*neighbourhood distances). We chose 50 m as the *ε-*neighbourhood distance for downstream analyses, as we estimate it to be the maximum distance at which coatis at our study sites would be able to communicate with one another to coordinate their movements. However, our results were qualitatively similar for *ε* values of 30 m and 70 m (see S7 – 12). For every 10-minute time step, we recorded the number and identities of individuals in each subgroup. Note that a subgroup by this definition may contain a single individual (if it is not within *ε* distance of any other group member).

One potential downside of this clustering method is that it can result in a likely-spurious “group merge” if two subgroups are between *ε* and 2x *ε* distance from one another and one individual moves between the two subgroups, momentarily joining them together. However, in our downsampled data, we did not observe any instances of this edge case.

#### Characterising subgrouping patterns

To determine the proportion of time groups were split into subgroups, we calculated the frequency distribution of the number of subgroups across the entire collaring period for each study group. To determine how subgroups were divided, we filtered these data to periods when the group was split into two and three subgroups and calculated the frequency distribution of the number of members in each subgroup.

We defined *group splits* as instances in which a group, at time *t*, split into two or more subgroups in the next time step *t*+1 (10 minutes later) (Figure 1). We excluded instances where a single individual (often the adult male) left the group or entered the group. For cases when individuals had missing data (due to poor satellite connection) before or after a split event, their data for that event was excluded from analyses (i.e. that individual was not considered part of any groups (see Figure S2 for details on missing data). We calculated the duration of splits as the time between the last split event to the next merge event, excluding single individuals. This was because there was the possibility of further splits occurring in a group that had previously split off from the full group.

**Figure 1.**
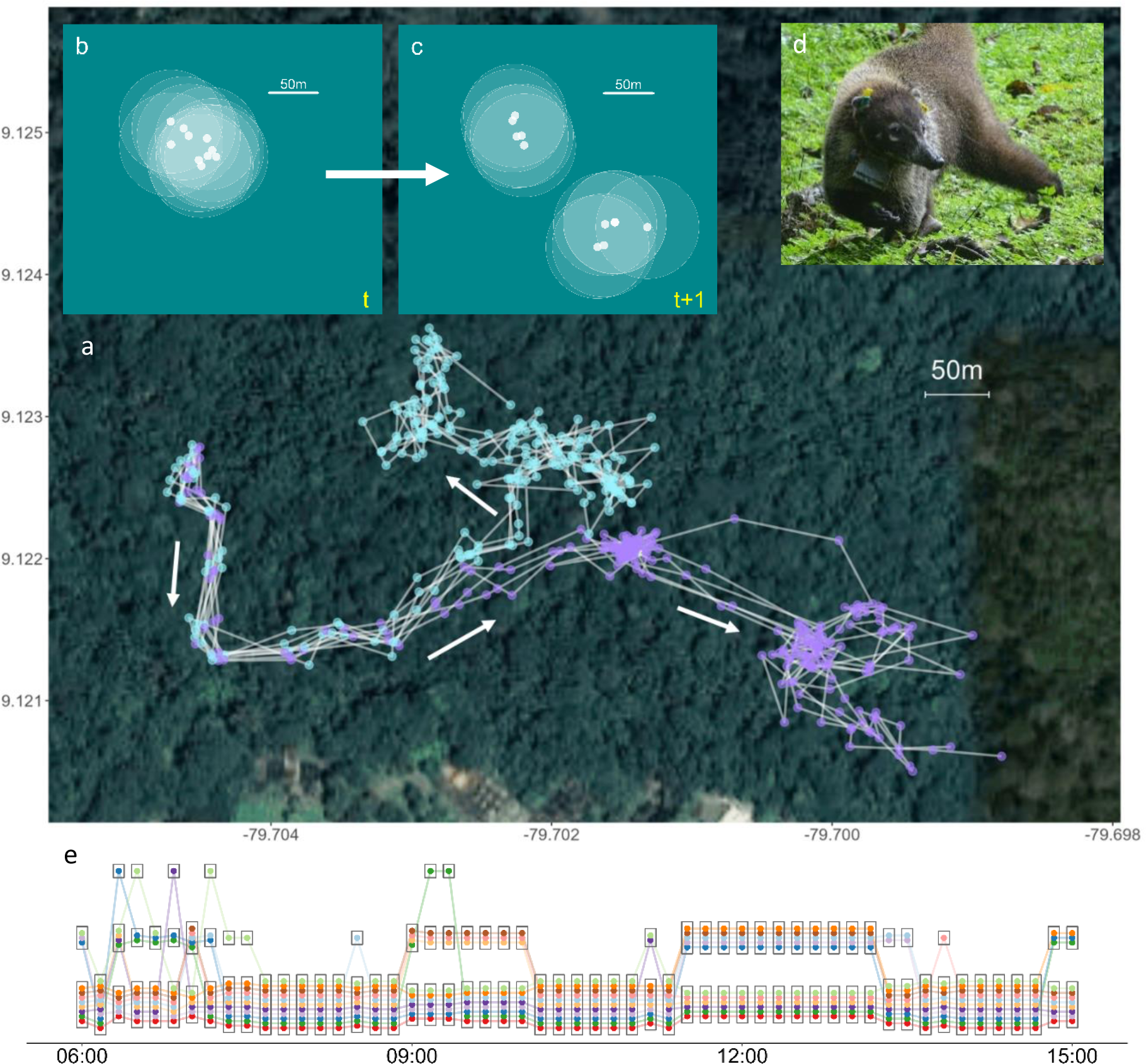
Whole-group tracking and quantifying fission-fusion dynamics. (a) Example of movement trajectories of coati group members during one day (0900 – 1800). Data are shown from the Galaxy group on the 28.12.21, excluding the single adult male. Points are coloured by the resultant subgroup, and arrows denote the direction of travel. (b-c) Illustration of how subgroups and group splits were defined from GPS data. (b) shows an initial moment in time (*t*) when the full group was together, and (c) shows a moment 10 minutes later (*t*+1) when the group had split into two subgroups. Outlined white circles represent the 50m *ε-*neighbourhood for each group member; overlapping groups of circles are considered distinct subgroups. (d) Photo of an adult female coati wearing a tracking collar, and ear tags used for identification. (e) Visualization of subgrouping patterns of a coati group over a 12-hour period, with x-axis indicating time and y-axis indicating subgroup membership. Coloured points represent individuals, and outlined groups of points represents subgroups identified at each moment in time.

#### Quantifying consistency of subgroup membership

To quantify subgrouping preferences, we first calculated the overall proportion of time each dyad was in the same subgroup for all days tracked. To account for missing data, we only incorporated times when both individuals’ locations were known into this calculation.

To determine whether subgrouping patterns were driven by one long fission event or from repeatedly splitting with the same individuals over time, we assessed whether subgroup membership across repeated group splits showed consistent patterns, i.e. whether certain pairs of individuals tended to be in the same subgroup at a rate greater than that expected by chance. Across all group splits (see above definition), we first computed the probability, 𝑝_𝑖𝑗_, that each pair of individuals 𝑖 and 𝑗 split into the same subgroup, given that both were present in the original group before the split occurred. We then defined the *consistency* of the subgrouping patterns across the entire group as

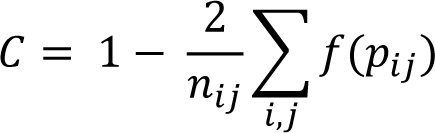

where 𝑛_𝑖𝑗_ is the total number of dyads and 𝑓(𝑝_𝑖𝑗_) is a function whose value equals 𝑝_𝑖𝑗_ if 𝑝_𝑖𝑗_ < 0.5 and 1 − 𝑝_𝑖𝑗_ otherwise. The value of 𝐶 ranges from 0 to 1, with the theoretical minimum of 0 occurring if all dyads have a probability of 50% of joining the same subgroup, and the theoretical maximum of 1 occurring if each dyad is either always (𝑝_𝑖𝑗_ = 0) or never (𝑝_𝑖𝑗_ = 1) in the same subgroup.

After computing the consistency *C* for the real data from each study group, we compared these values to permuted data assuming that individuals joined subgroups at random during each group split. To construct this null model, we maintained the same number of real splits as occurred in the data as well as the same subgroup sizes and individuals involved in each split, but randomly allocated individuals to the different subgroups for each split. We repeated this procedure 1000 times, computing the value of *C* for each artificially randomised dataset and generated a distribution of these expected *C* values under the assumption of random subgroup assignment. Finally, we compared this null distribution to the value of *C* observed in the real data to determine how likely the real or a greater level of consistency would be to occur based on the random null model (i.e. to compute a p-value).

#### Quantifying the role of social and physiological factors with subgroup membership

We used a Multiple Regression Quadratic Assignment Procedure (MRQAP) with the Double Semipartialling (DSP) method to determine whether age homophily (i.e. individuals of the same age), sex homophily, or relatedness was associated with subgroup membership (Dekker et al., 2007). The dependent matrix was the proportion of time each dyad was in the same subgroup when the group had split and the independent matrices were age-based homophily, sex-based homophily, and genetic relatedness. When dyads in the homophily matrices were the same, they were given a value of 1, while dissimilar dyads were given a value of 0. All networks were undirected.

## Results

The three study groups differed in their fission-fusion dynamics: two (Galaxy and Presidente) often split into multiple subgroups, whereas the third (Trago) remained cohesive and did not exhibit fission-fusion dynamics (Figure S3). The third group was therefore excluded from further analyses of subgrouping dynamics. The frequency of fission events and the duration of time that groups remained split into subgroups differed between the groups: over the tracking period, Galaxy group split 29 times whereas Presidente group split 43 times, with median split durations of 10.5h (IQR = 2.7h - 15.3h) and 2h (IQR = 1.3h – 4.3h) respectively. We found no consistent pattern in the time of day when fissions occurred for either group (Figure S4). The two groups also varied in the size of subgroups formed during fission events. Galaxy group tended to remain cohesive or split into either two or three subgroups, with the most common number of subgroups being two (Figure 2a). When there were two subgroups, the group split either evenly or the majority of the group were together and one individual was alone (Figure 2b). A similar pattern was observed when there were three subgroups, which typically occurred when one individual was alone and the rest of the group was evenly split (Figure 2c). Presidente group exhibited similar subgrouping patterns (Figure 2d), but subgroup sizes were often unevenly divided into a larger group of ∼12 individuals and a smaller group of ∼4 individuals (Figure 2e). When divided into three groups, subgroups of a range of sizes were observed (Figure 2f).

**Figure 2.**
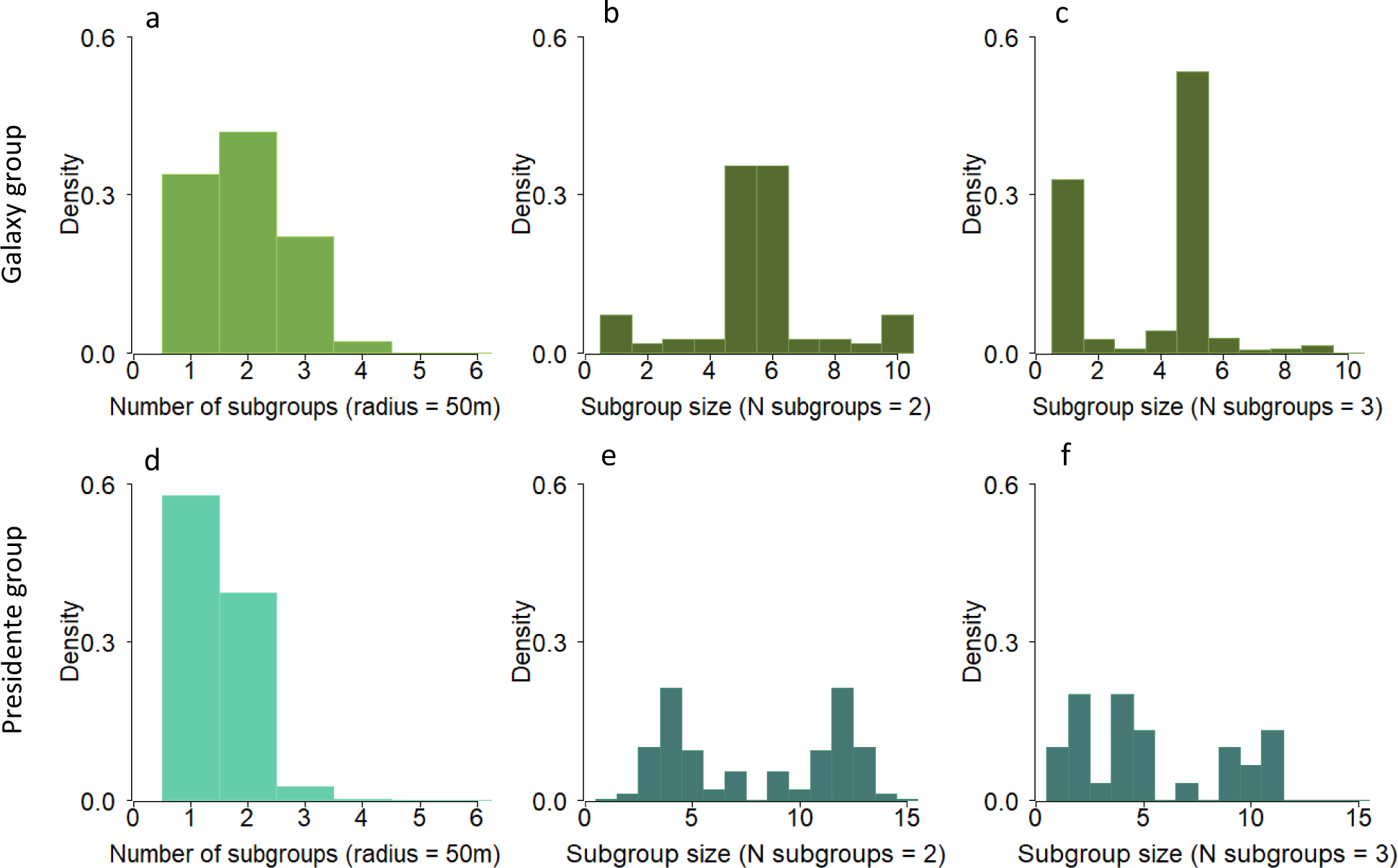
Number of subgroups and subgroup sizes across two coati groups. (a, d) Probability of observing different numbers of subgroups at any moment in time for two different coati groups. (b-f) Histograms of the number of individuals in each subgroup when the group was split into two (b, e) or three subgroups (c, f). Top row (a-c) shows data from the Galaxy group and bottom row (d-f) shows data from the Presidente group. The Trago group remained together for the entire duration of the tracking period (i.e. one subgroup), so data are not shown here.

Coatis showed high consistency in subgroup membership across splitting events (P < .001 for both groups based on permutation tests, Figure S5). When not together, both groups were most often split into two specific subgroups (Figure 3a, c, yellow blocks). In Galaxy group, one of the subgroups was composed of four adult females and one juvenile, while the other subgroup consisted of all three subadults and two adult females. The single adult male was more often alone compared to group members. When he was with one of the subgroups, however, he tended to associate with the adult females. Presidente group tended to split unevenly, with the smaller subgroup composed of two subadults and one adult female and the larger subgroup composed of eight juveniles, three subadults and two adult females. In both study groups, relatedness was a significant predictor of subgroup membership (Table 2; Figure 3b, d; Figure S6). In contrast, age and sex homophily showed no significant effect on subgroup membership in either group.

**Figure 3.**
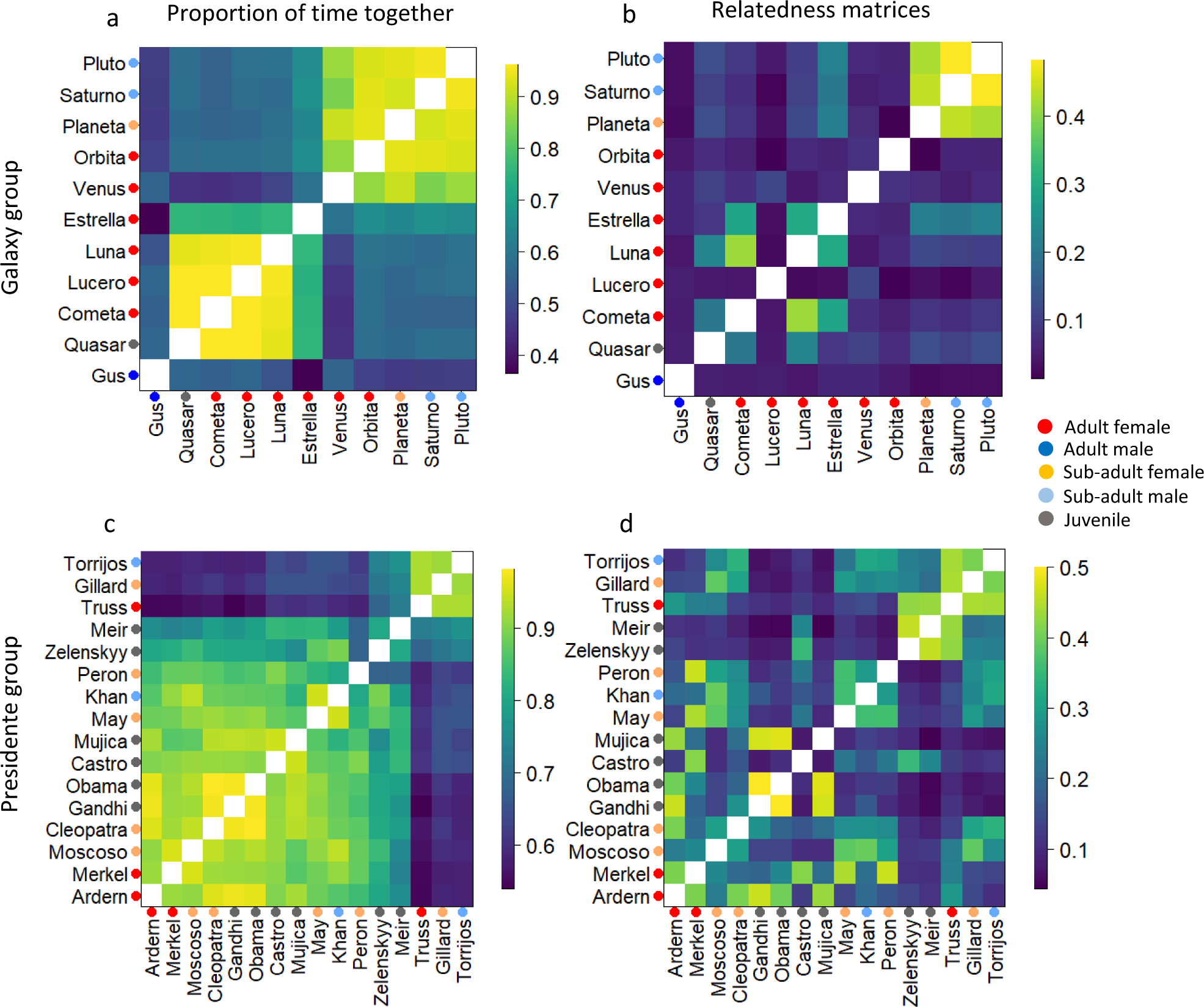
Two consistent subgroups were observed in the coati groups that exhibited fission-fusion dynamics. Left panels (a and c) show association matrices representing the proportion of time each dyad was in the same subgroup across the full dataset for Galaxy (a) and Presidente (c) groups respectively. Rows and columns of each matrix represent individuals, with coloured points representing age/sex class of each individual. Coloured squares in each matrix indicate the proportion of time each dyad was found in the same subgroup, across all times when both individuals in the dyad were tracked. Right panels (b and d) show the relatedness matrices for Galaxy group (b), Presidente group (d), using the Triadic Maximum Likelihood method. This estimator has coefficients of relatedness ranging from 0 to 1 (full siblings and offspring are approximately 0.5, half siblings are 0.25, and aunts and uncles are 0.125).

**Table 2.** Results of MRQAP regression predicting subgroup membership based on relatedness, age homophily, and sex homophily between dyads for both groups. The dependent matrix is subgroup membership, defined as the proportion of time each dyad was found in the same subgroup across the full dataset. Independent matrices are age and sex homophily (1 if dyad were in the same class, 0 if dyad were in a different class).

| Group |  | Galaxy group |  |  | Presidente group |  |  |
| --- | --- | --- | --- | --- | --- | --- | --- |
| Dependent matrix | Independent matrices | Coefficient | P | Adjusted R<sup>2</sup> | Coefficient | P | Adjusted R<sup>2</sup> |
| Subgroup membership | Age | -0.041 | 0.399 | 0.169 | -0.030 | 0.235 | 0.085 |
|  | Sex | 0.059 | 0.247 |  | -0.025 | 0.207 |  |
|  | Relatedness | 0.710 | **0.001** |  | 0.363 | **0.003** |  |

## Discussion

Whole-group GPS tracking revealed substantial variation in the extent and nature of fission-fusion dynamics of coatis, while also demonstrating common drivers of subgroup membership. In alignment with findings from captive populations (Romero & Aureli, 2007), we found that coatis repeatedly formed subgroups with the same set of individuals and that these subgroups tended to consist of related individuals rather than individuals of the same age or sex. Genetic relatedness has been shown to influence patterns of association in a variety of species that show fission-fusion dynamics, including spotted hyenas (Holekamp et al., 1997, 2012; Van Horn et al., 2004), African elephants (Archie et al., 2006), and orangutans (van Noordwijk et al., 2012). A common feature across many of these animal societies is the integration of individuals with varying levels of relatedness (e.g. multiple matrilines) into the same social group or community, as well as the maintenance of long-term, differentiated social relationships among group members that are often correlated with relatedness (Archie et al., 2006; Carter et al., 2013; Sueur et al., 2010). Within such complex social landscapes, consistent subgrouping patterns may result from a preference to maintain cohesion with specific group members who provide social benefits, by avoiding individuals who are costly to be near, or a combination of both mechanisms (Romero & Aureli, 2007, 2008). In coatis, individuals tend to have stronger affiliative relationships with their close relatives, which is often exhibited by grooming and coalitionary support during aggressive encounters (Hirsch et al., 2012). Such close relationships have been shown to provide major fitness benefits in other species (Silk, 2007), hence maintaining cohesion with related individuals is likely to come with fitness benefits in coatis. On the other hand, forming subgroups with related individuals might be a strategy to avoid unrelated group members. Previous studies have shown that coatis not related to other group members receive far more aggression (Gompper et al., 1997). Distancing themselves from unrelated individuals could be an adaptive strategy for individuals to minimise their risk of receiving aggression during foraging. Further work investigating the behavioural context and social interactions that occur before, during, and after group splits could help differentiate between these possible underlying mechanisms.

Relatedness-driven subgrouping may have important implications for the processes of collective decision-making in social groups. Consensus costs based on differing preferences are commonly invoked to explain patterns of group cohesion and decision-making across species, with group splits being an alternative to achieving consensus (Conradt & Roper, 2005). If individuals split with related group members, however, group fissions may not produce subgroups whose preferences are more aligned than the group as a whole. With high potential consensus costs in such subgroups, mechanisms of decision-making and the distribution of influence over collective decisions will have an important effect on the costs of grouping. Investigating the extent to which coordination among group members plays a role in subgroup formation and subsequent fission events will provide further insights into the decision-making mechanisms within animal groups.

Our results do not support physiological drivers as being the primary determinant of subgrouping patterns in coatis. These results contrast with patterns seen in several other fission-fusion species, where physiological differences play a role in subgrouping patterns (Bond et al., 2019; Galezo et al., 2018; Hunt et al., 2019; Surbeck et al., 2017). In spider monkeys, social, ecological, and physiological factors influence their fission-fusion dynamics (Aguilar-Melo et al., 2018; Hartwell et al., 2021), and previous studies have found that group members often split with individuals of similar nutritional requirements, which correlates with sex homophily (Hartwell et al., 2014; Rodrigues, 2014). The lack of support for physiological drivers of group splits could reflect a lack of strong differences in preferred foraging patches or travel speed in this species. Alternatively, such differences may exist but are not strong enough to outweigh social drivers, leading individuals to compromise their own physiological needs to remain associated with related individuals. Even though we did not find any effect of age and sex homophily on subgrouping patterns, other physiological factors which are not associated with these broad categories could still play a role in driving subgroup composition.

Although we focused our investigation on the drivers of group splits, one of the groups we tracked did not split at all. This group (Trago) was composed of seven juveniles and one subadult male that was visibly limping. The other adult females and sub-adults had left the group to give birth or disperse before or shortly after collaring (sub-adult males normally leave their groups during this period). Juveniles are significantly smaller than adults and have a greater risk of predation compared to other group members (Hass & Valenzuela, 2002). The juveniles in this group might have remained more cohesive with one another to minimise their risk of predation, supporting a role of predation risk in shaping subgrouping patterns. This could have been particularly important given their relatively small group size (N=7). Alternately, juvenile cohesiveness could have been driven by social attraction, as juvenile coatis often spend a considerable amount of time playing and closely associate with one another within groups (Hirsch, 2011b, 2011a; Kaufmann, 1962). The remaining members of the Trago group were mostly closely related (Figure S3b), so the tendency to associate together might also reflect cohesion based on social relationships as seen in the other two groups.

Even though we identified some shared features in the subgrouping behaviour of coatis, our results also highlight the substantial variation in fission-fusion dynamics that can occur within the same species. The tendency to split, the durations of splits, and the relative subgroup sizes all varied across the three groups we studied. Differences in group size, demographic composition, and the relatedness of group members (Lehmann & Boesch, 2004) may all have contributed to this variation, highlighting the importance of studying multiple groups. The drivers of fission-fusion dynamics are likely to reflect a complex balance of factors that may fundamentally change depending on both the social and ecological context. Such variation can have important consequences for group members’ foraging efficiency and predation risk, ultimately impacting reproductive success and survival (Rubenstein, 1978). Collecting data on a larger sample of social groups under different ecological conditions could shed further light on the drivers of this variation. Additionally, expanding the approach we applied here to other study systems could yield comparable data across a range of species, enabling a broader investigation of the social, physiological, and ecological underpinnings of fission-fusion dynamics in animal societies.

## Acknowledgements

We thank the Smithsonian Tropical Research Institute for permission to conduct this research. Thank you to MiAmbiente and the Republic of Panama for the permission to export the biological samples. We thank Lil Camacho for processing the export permits for the biological samples. Thanks to Patrick Paetzold for helping design the 3D printed cases for the housing of the audio recorders. We thank Carolina Mitre Ramos, Brandol Ortega, and Lucía Torrez for helping with the capturing, collaring, and data collection on the coatis in Gamboa and Barro Colorado Island. Thank you to Rachel Page and Melissa Cano for their support during fieldwork. We also thank the members of the Communication and Collective Movement group as well as members of the Communication and Coordination Across Scales team for feedback and useful discussion. Finally, thank you to the members of the department for the Ecology of Animal Societies for fruitful discussion.

## Author Contributions

BTH, MCC, and ASP conceived the coati collaring study and secured the funding. EMG designed and tested the collars. EMG, JO, and BTH collected the tracking data. EMG, BTH, MCC, and ASP conceived the specific questions addressed in this study. ASP and EMG designed the analysis and wrote the analysis code. PM conduced a code review to ensure the accuracy and reproducibility of analysis. MJQ and EMG analysed the genetics data. EMG wrote the initial manuscript draft, and all authors contributed to, and approved, the final manuscript.

## Supplementary figures

**Figure S1.**
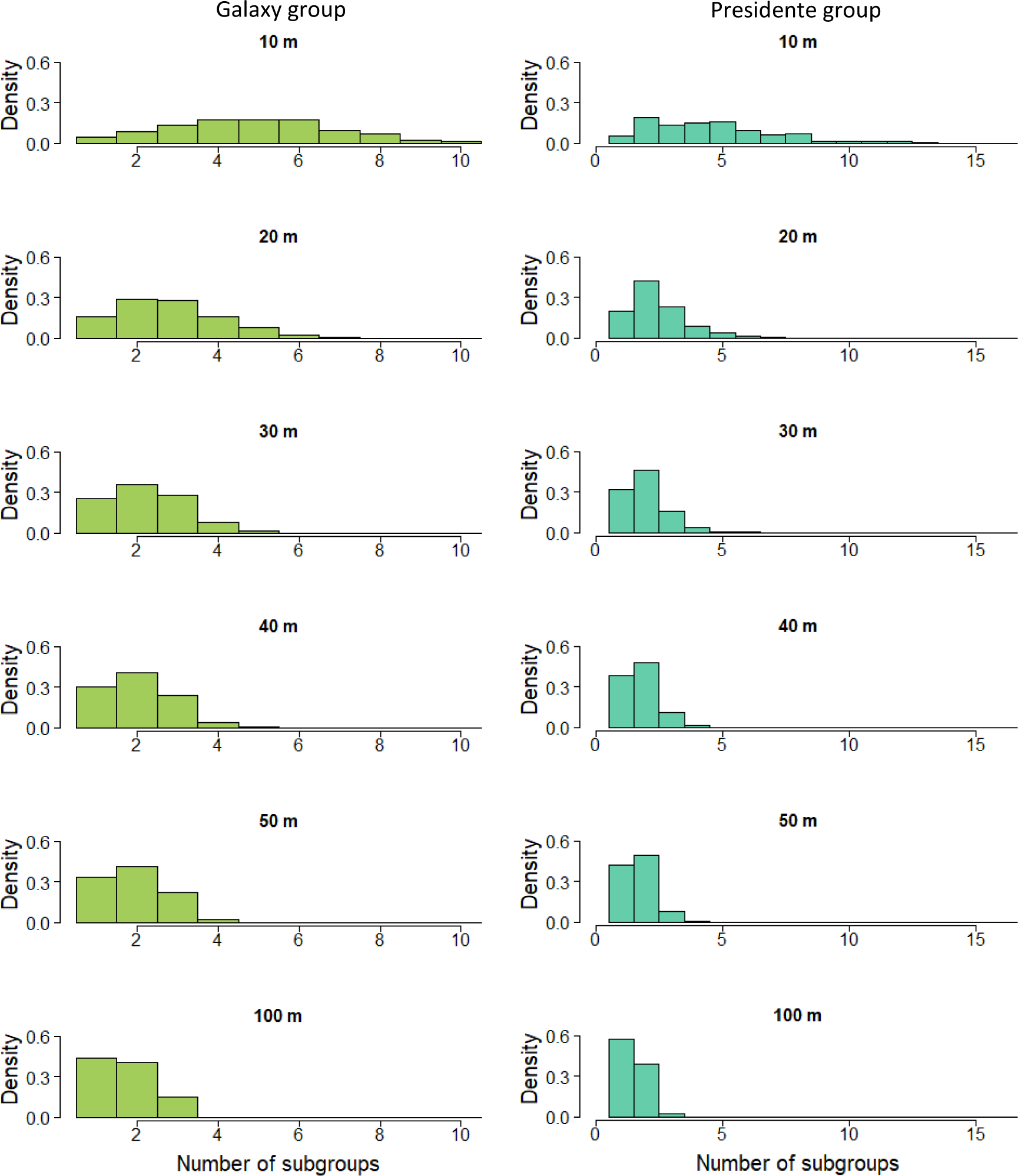
Number of subgroups for Galaxy group (left) and Presidente group (right) when the *ε-* neighbourhood distance is set to 10m, 20, 30m, 40m, 50m and 100m respectively.

**Figure S2.**
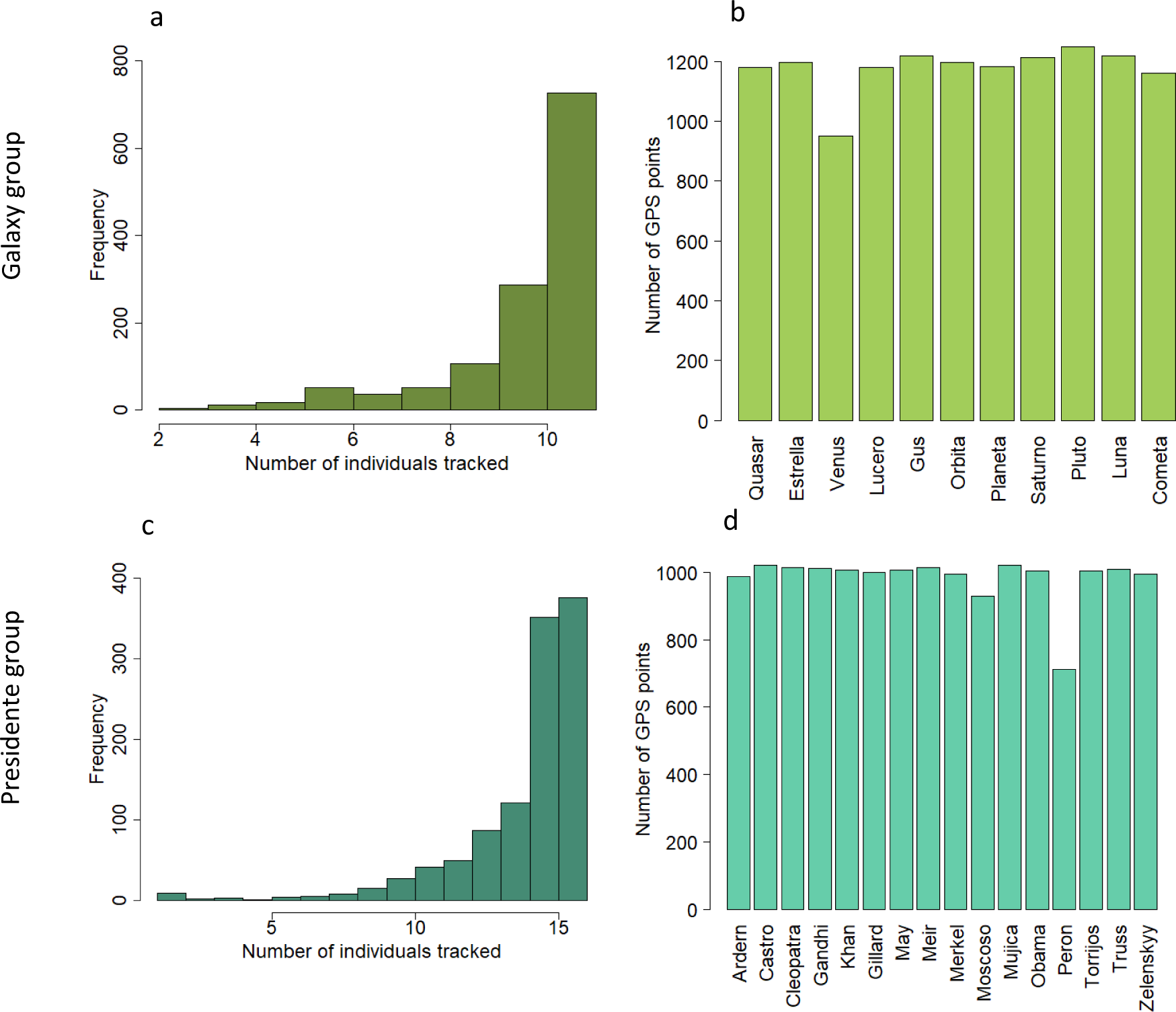
Histogram for number of individuals tracked via GPS during daytime hours for Galaxy group (a), and Presidente group (c). Bar plot for number of total GPS points from each individual used in this study’s analysis for Galaxy group (b), and Presidente group (d).

**Figure S3.**
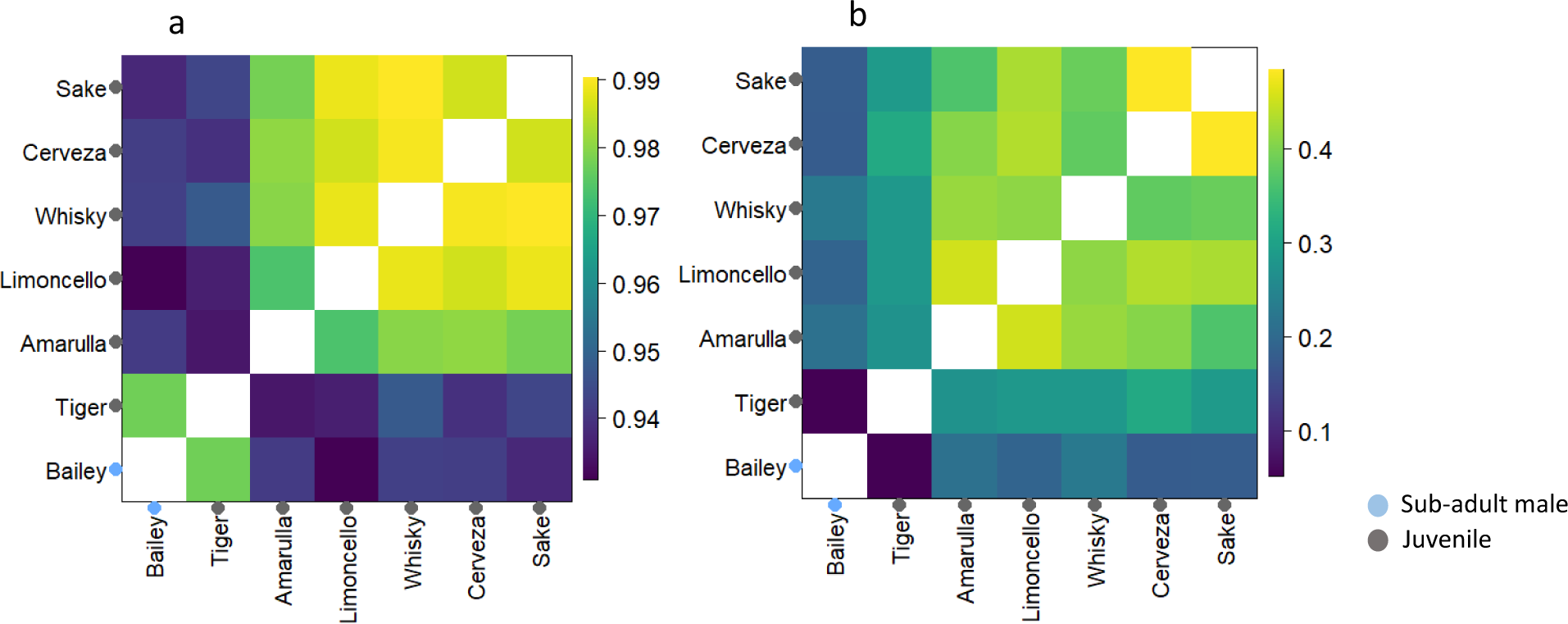
Proportion of time every each were in the same subgroup for Trago group (a) (measured using dbscan with 50m *ε-*neighbourhood distance), across all data where all group members had a GPS fix. Relatedness matrix for Trago group (b) using the Triadic Maximum Likelihood method. This estimator has coefficients of relatedness ranging from 0 to 1 (full siblings and offspring are approximately 0.5, half siblings are 0.25, and aunts and uncles are 0.125).

**Figure S4.**
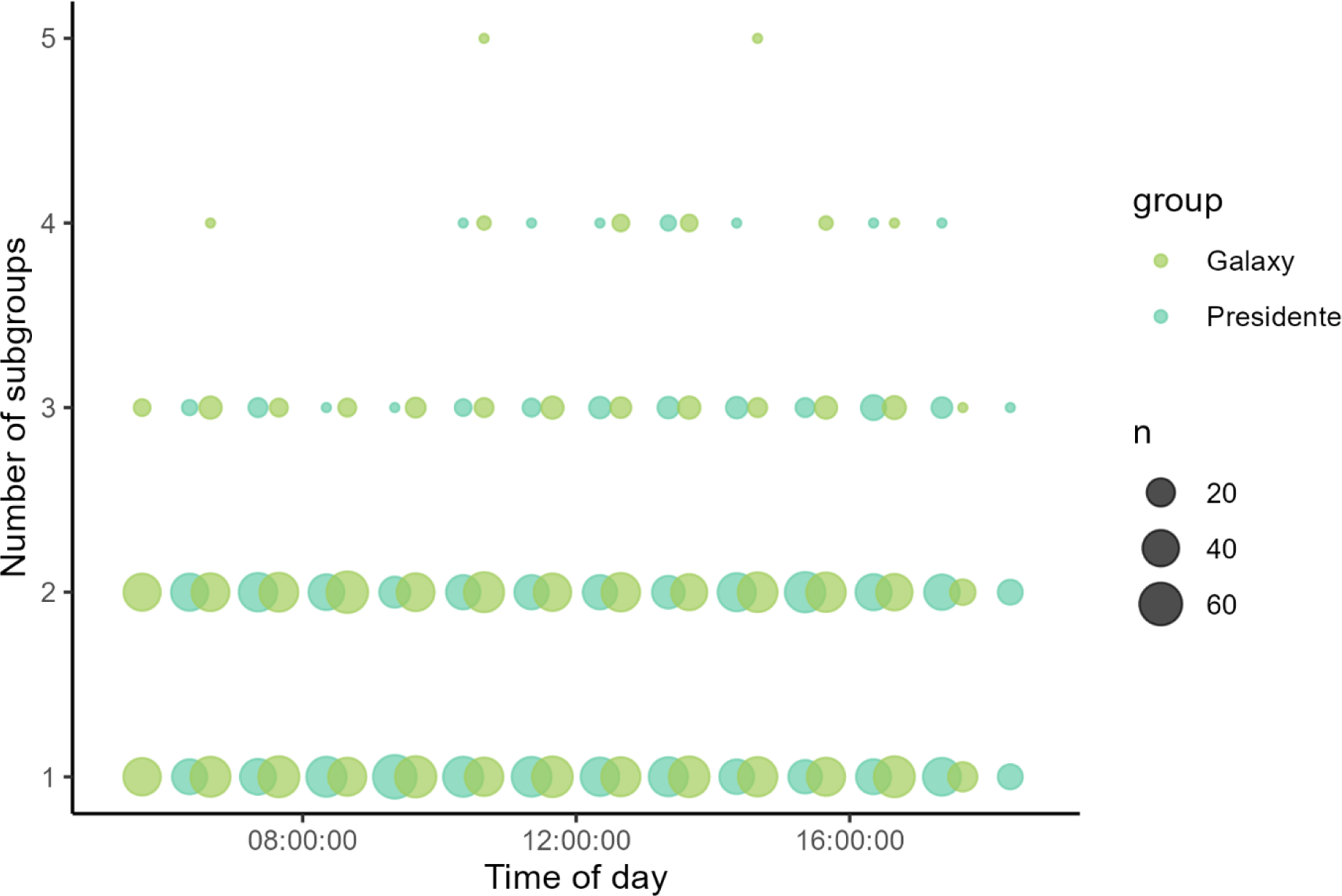
Number of subgroups for each hour of the day for Galaxy group and Presidente group. The size of the point is proportional to the frequency of data points for each time of day across the total collar period for both groups.

**Figure S5.**
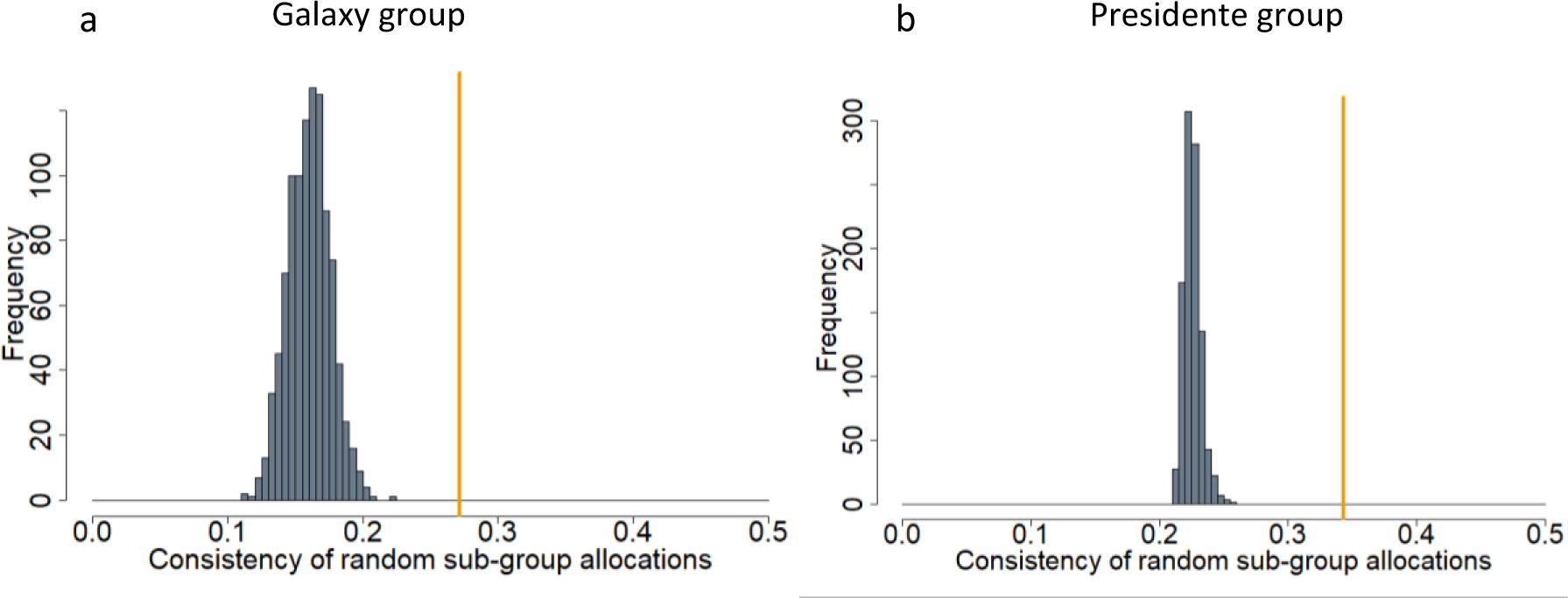
Consistent subgroup membership across group splits. Histograms show the distribution of the subgroup consistency metric under a null model assuming random allocations of group members to subgroups during group splits (1000 permutations) for Galaxy group (a), and Presidente group (b). Orange vertical lines show the consistency value for the real split data.

**Figure S6.**
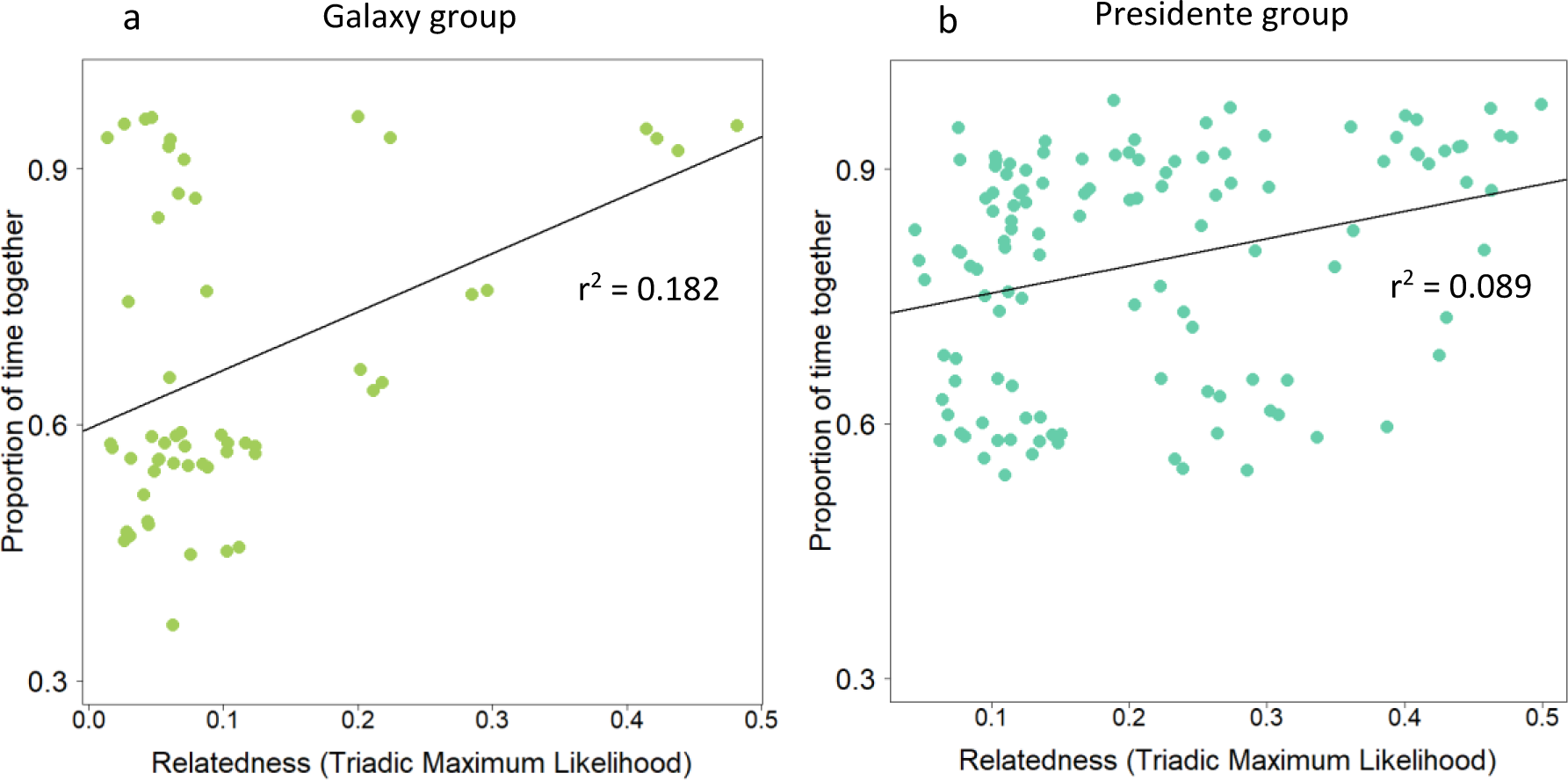
Positive correlation between relatedness and proportion of time together (subgroup membership matrices) for Galaxy group (a), and Presidente group (b). Black line is the linear regression and the r^2^ value is denoted.

### Alternative parameterisations

To test the robustness of our results to the distance threshold chosen for determining subgroups (*ε* = 50m), we re-ran the analyses in the main text with alternative thresholds of *ε* = 30m and *ε =* 70m. The results are shown in Figures S7-12 and Tables S1-2, which are comparable to the main analyses represented by Figures 2-3, Figure S5, and Table 1. Results did not differ qualitatively between these three different parameterizations.

**Figure S7.**
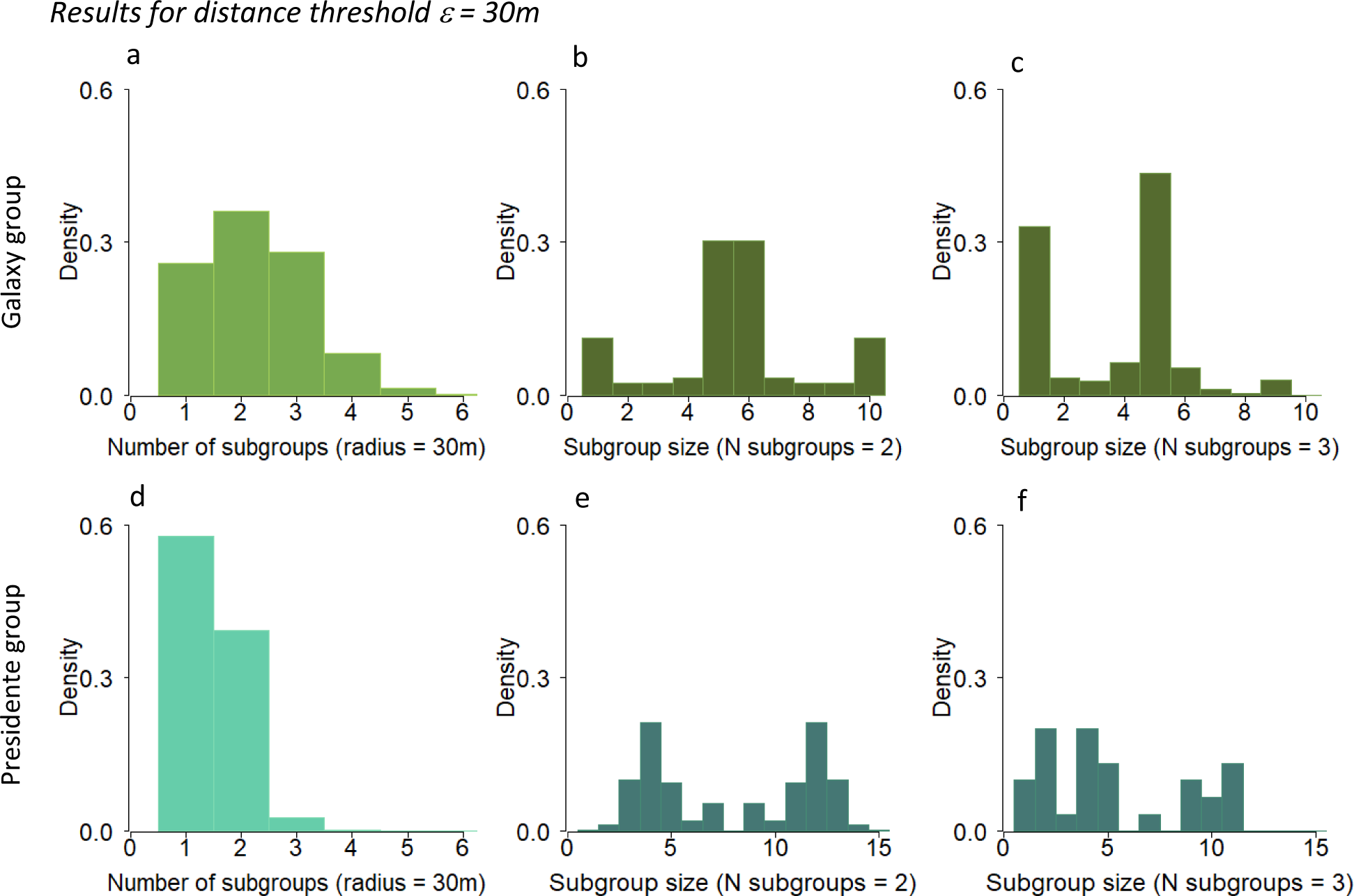
Characterisation of subgrouping patterns for Galaxy group (a, b, and c) and Presidente group (d, e, and f) when the *ε*-neighbourhood distance was set to 30m (compare to main text Figure 2). (a, d) Histogram of the probability of finding subgroups of different sizes in the tracking data. (b, e) Histograms of the number of individuals in each subgroup when the group was split into two subgroups, and three subgroups (c, f).

**Figure S8.**
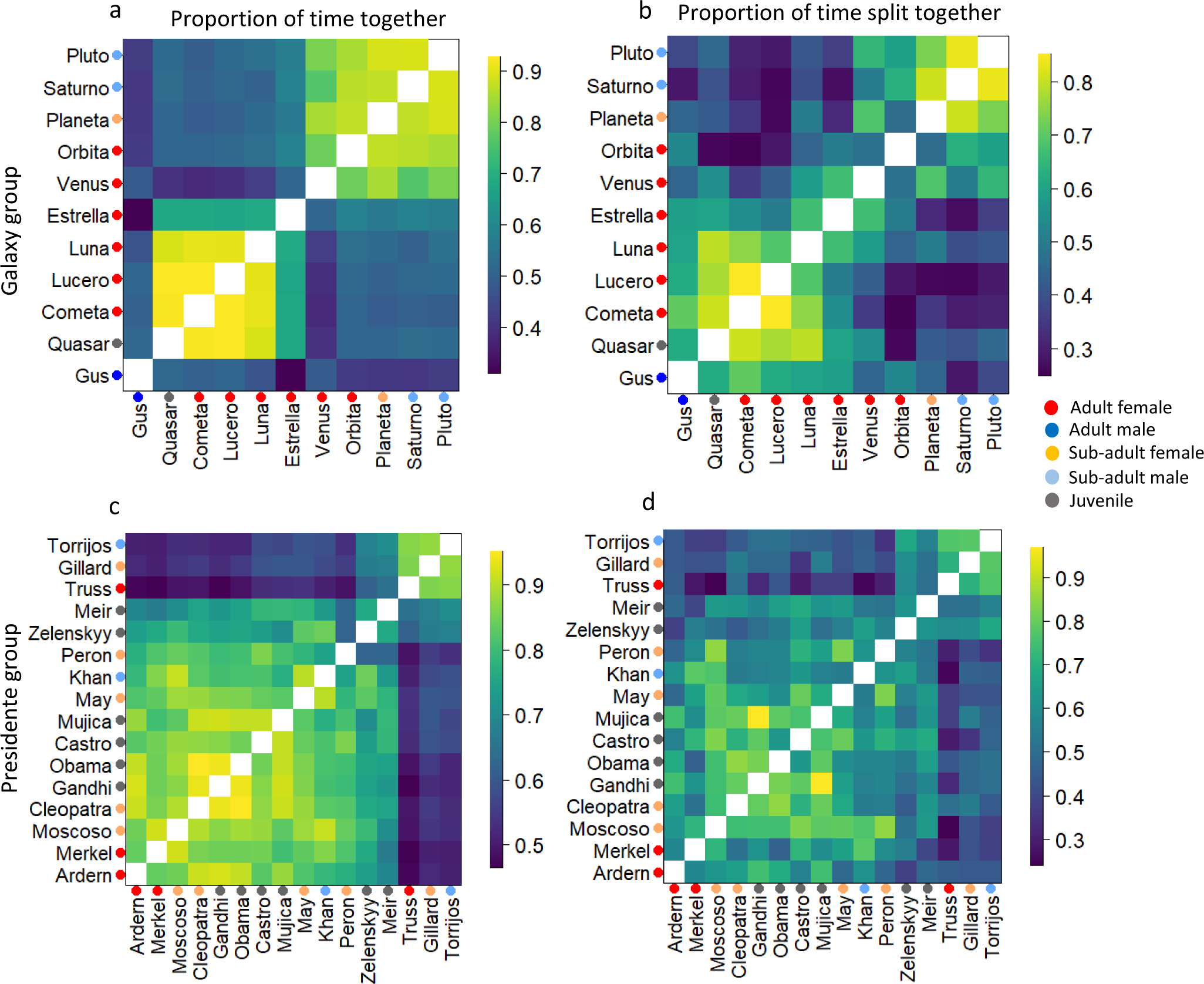
Two distinct subgroups were observed in the coati groups that exhibited fission-fusion dynamics. Results are shown for the parameterization *ε* = 30m (compare to main text Figure 3). Subgrouping patterns in Galaxy group (a,b) and Presidente group (c,d). Left panels (a and c) show association matrices representing the proportion of time each dyad was in the same subgroup across the full dataset for Galaxy (a) and Presidente (c) groups respectively. Row and columns of each matrix represent individuals, with coloured points representing age/sex class of each individual. Coloured squares in each matrix indicate the proportion of time each dyad was found in the same subgroup, across all times when both individuals in the dyad were tracked. Right panels (b and d) show the proportion of time individuals in each dyad joined the same subgroup during events when the full group split into subgroups, for Galaxy (b) and Presidente (d) groups respectively.

**Figure S9.**
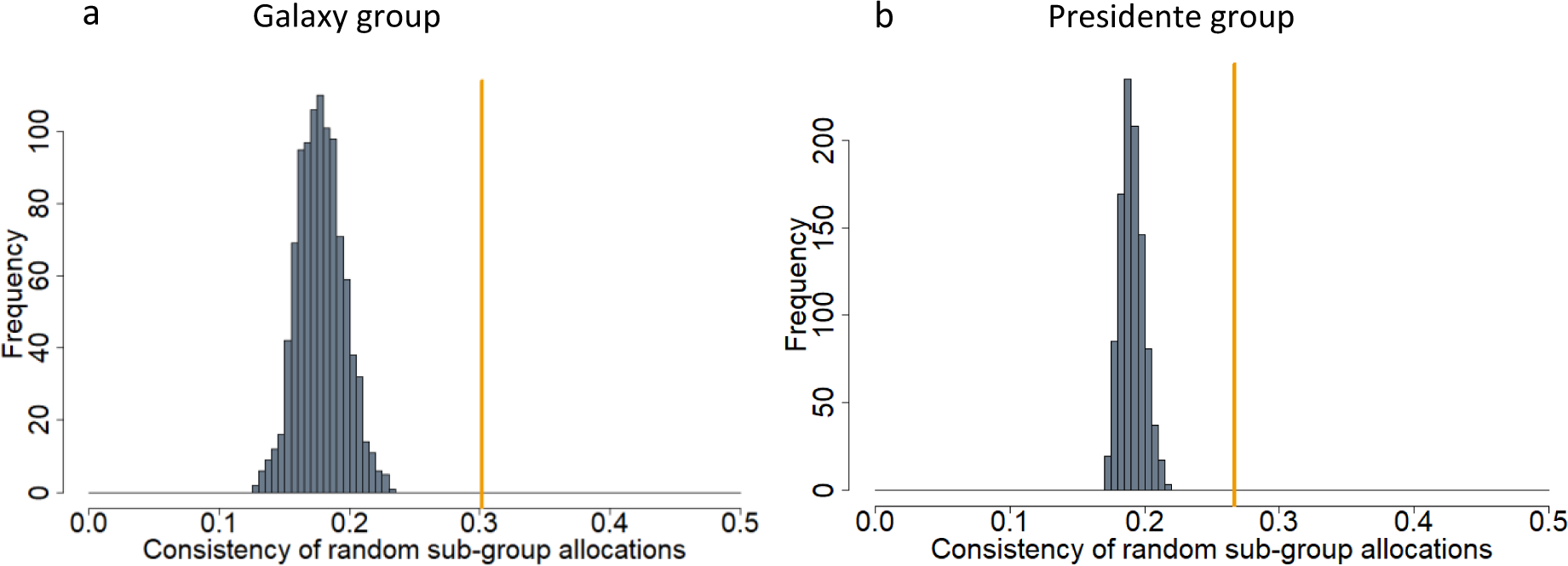
Consistent subgroup membership across group splits. Results are shown for the parameterization *ε* = 30m (compare to main analysis shown in Figure S5). Histograms show the distribution of the subgroup consistency metric under a null model assuming random allocations of group members to subgroups during group splits (1000 permutations) for Galaxy group (a), and Presidente group (b). Orange line show the consistency value for the real split data.

**Figure S10.**
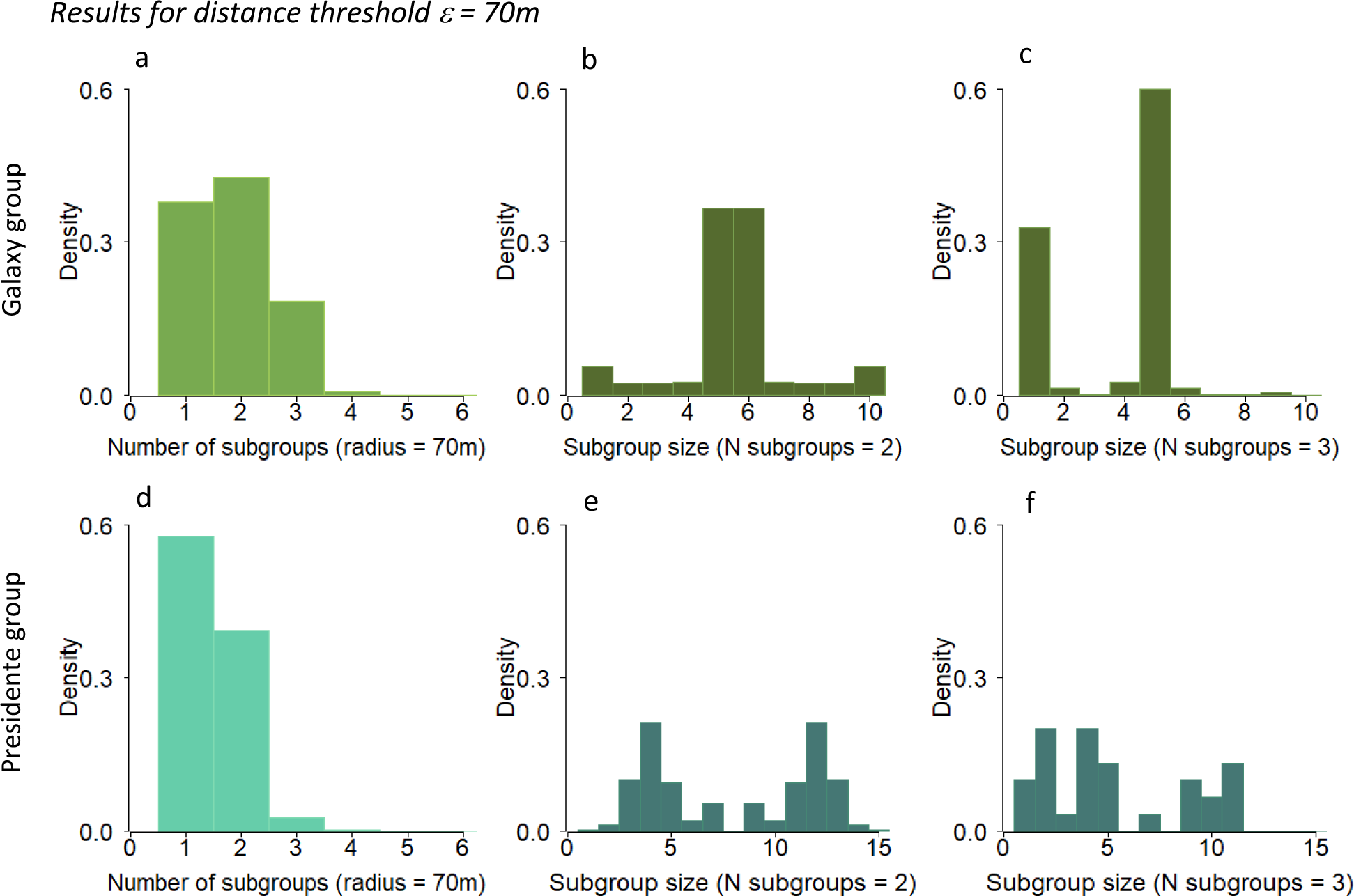
Characterisation of subgrouping patterns for Galaxy group (a, b, and c) and Presidente group (d, e, and f) when the *ε-*neighbourhood distance was 70m (compare to main text Figure 2). (a, d) Histogram of the probability of finding subgroups of different sizes in the tracking data. (b, e) Histograms of the number of individuals in each subgroup when the group was split into two subgroups, and three subgroups (c, f).

**Figure S11.**
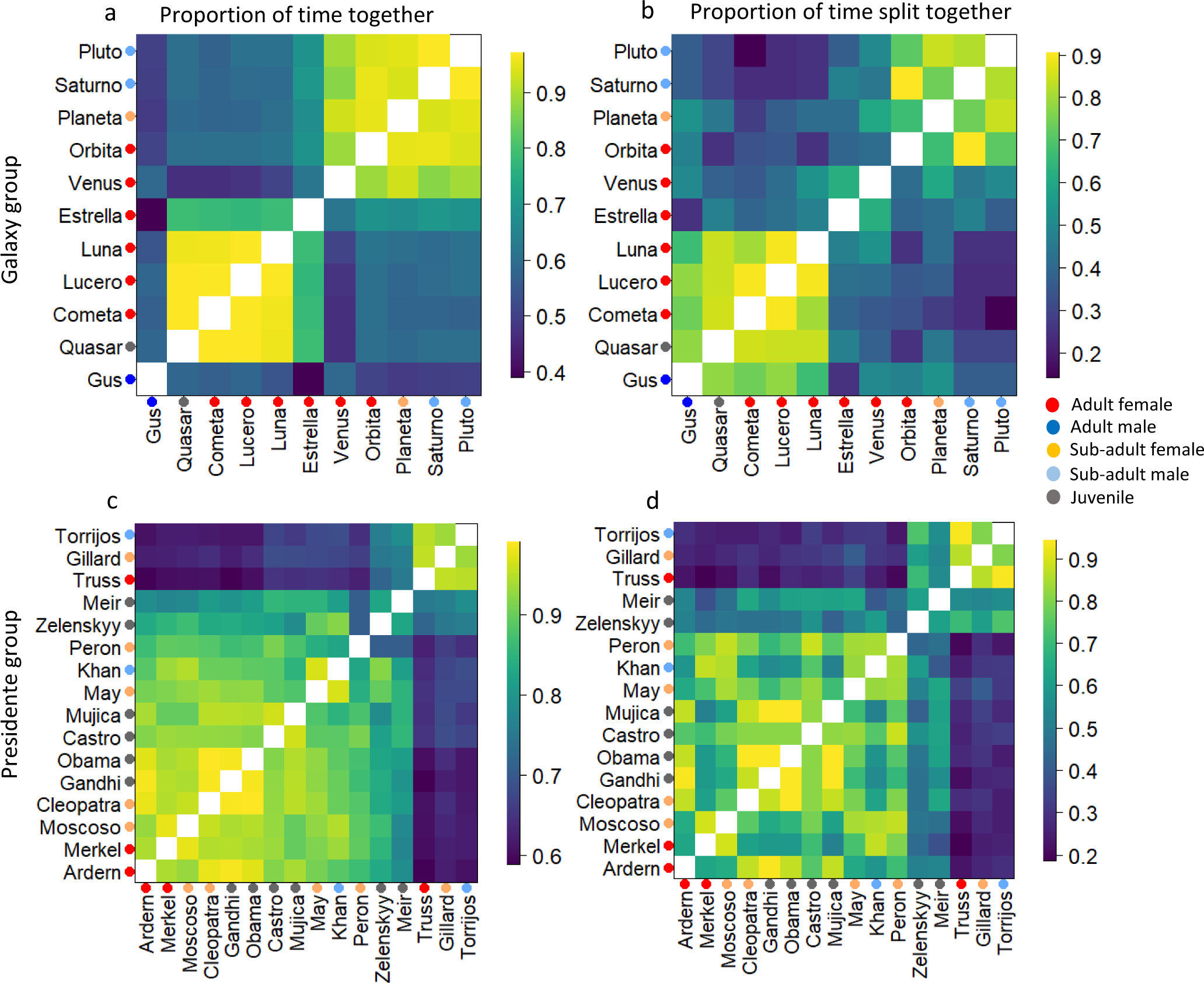
Two distinct subgroups were observed in the coati groups that exhibited fission-fusion dynamics. Results are shown for the parameterization *ε* = 70m (compare to main text Figure 3). Subgrouping patterns in Galaxy group (a,b) and Presidente group (c,d). Left panels (a and c) show association matrices representing the proportion of time each dyad was in the same subgroup across the full dataset for Galaxy (a) and Presidente (c) groups respectively. Row and columns of each matrix represent individuals, with coloured points representing age/sex class of each individual. Coloured squares in each matrix indicate the proportion of time each dyad was found in the same subgroup, across all times when both individuals in the dyad were tracked. Right panels (b and d) show the proportion of time individuals in each dyad joined the same subgroup during events when the full group split into subgroups, for Galaxy (b) and Presidente (d) groups respectively.

**Figure S12.**
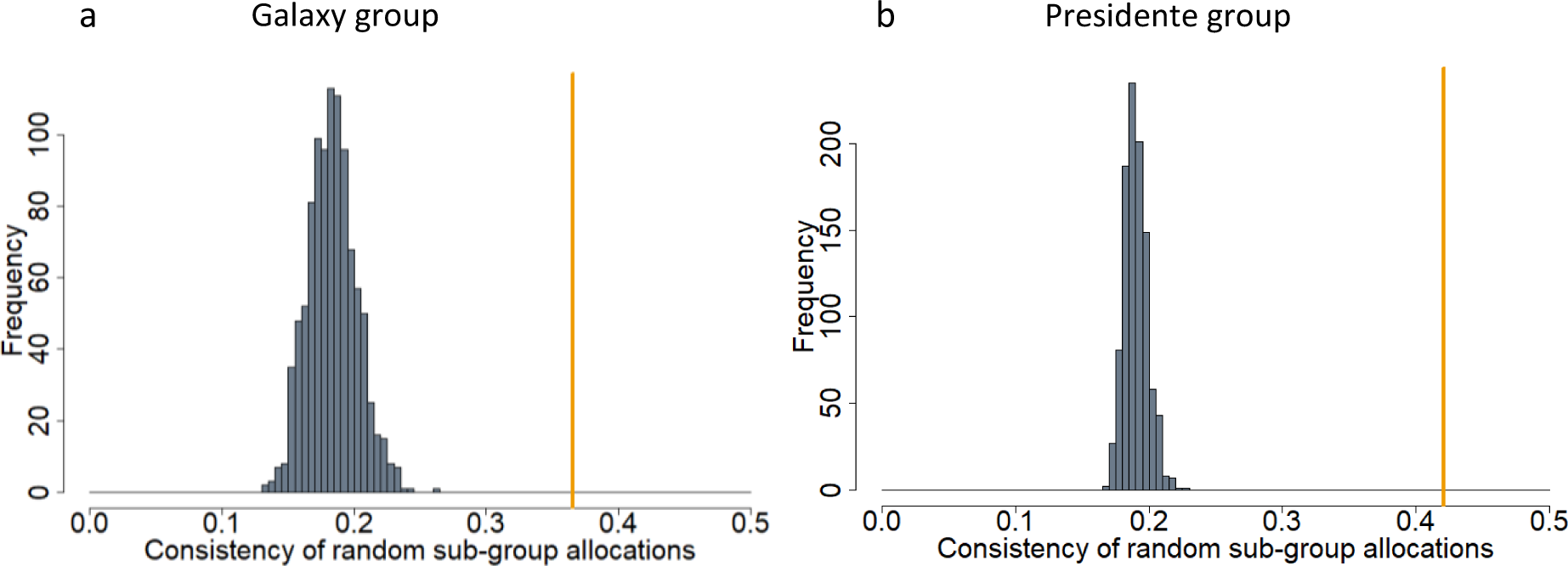
Consistent subgroup membership across group splits. Results are shown for the parameterization *ε* = 30m (compare to main analysis shown in Figure S5). Histograms show the distribution of the subgroup consistency metric under a null model assuming random allocations of group members to subgroups during group splits (1000 permutations) for Galaxy group (a), and Presidente group (b). Orange line show the consistency value for the real split data.

**Table S1.** Results of MRQAP regression predicting subgroup membership based on relatedness, age homophily, and sex homophily between dyads for both groups. Results are shown for the parameterization *ε* = 30m (compare to main text Table 1). The dependent matrix is subgroup membership, defined as the proportion of time each dyad was found in the same subgroup across the total dataset. Independent matrices are age and sex homophily (1 if dyad were in the same class, 0 if dyad were in a different class).

| Group |  | Galaxy group |  |  | Presidente group |  |  |
| --- | --- | --- | --- | --- | --- | --- | --- |
| Dependent matrix | Independent matrices | Coefficient | P | Adjusted R<sup>2</sup> | Coefficient | P | Adjusted R<sup>2</sup> |
| Subgroup membership | Age | -0.040 | 0.409 |  | -0.032 | 0.161 |  |
|  | Sex | 0.058 | 0.262 | **0.168** | -0.022 | 0.230 | 0.086 |
|  | Relatedness | 0.715 | **0.002** |  | 0.345 | **0.003** |  |

**Table S2.** Results of MRQAP regression predicting subgroup membership based on relatedness, age homophily, and sex homophily between dyads for both groups. Results are shown for the parameterization *ε* = 70m (compare to main text Table 1). The dependent matrix is subgroup membership, defined as the proportion of time each dyad was found in the same subgroup across the total dataset. Independent matrices are age and sex homophily (1 if dyad were in the same class, 0 if dyad were in a different class).

| Group |  | Galaxy group |  |  | Presidente group |  |  |
| --- | --- | --- | --- | --- | --- | --- | --- |
| Dependent matrix | Independent matrices | Coefficient | P | Adjusted R<sup>2</sup> | Coefficient | P | Adjusted R<sup>2</sup> |
| Subgroup membership | Age | -0.041 | 0.389 | **0.172** | -0.032 | 0.162 | 0.086 |
|  | Sex | 0.058 | 0.236 |  | -0.022 | 0.230 |  |
|  | Relatedness | 0.701 | **0.001** |  | 0.345 | **0.003** |  |

